# Normative Decision Rules in Changing Environments

**DOI:** 10.1101/2022.04.27.489722

**Authors:** Nicholas W Barendregt, Joshua I Gold, Krešimir Josić, Zachary P Kilpatrick

## Abstract

Models based on normative principles have played a major role in our understanding of how the brain forms decisions. However, these models have typically been derived for simple, stable environments, and their relevance to decisions under more naturalistic, dynamic conditions is unclear. We previously derived a normative decision model in which evidence accumulation is adapted to environmental dynamics (*Glaze et al., 2015*), but the evolution of commitment rules (e.g., thresholds on the accumulated evidence) under such dynamic conditions is not fully understood. Here we derive a normative model for decisions based on changing evidence or reward. In these cases, performance (reward rate) is maximized using adaptive decision thresholds that best account for diverse environmental changes, in contrast to predictions of many previous decision models. These adaptive thresholds exhibit several distinct temporal motifs that depend on the specific, predicted and experienced changes in task conditions. These adaptive decision strategies perform robustly even when implemented imperfectly (noisily) and can account for observed response times on a task with time-varying evidence better than commonly used constant-threshold or urgency-gating models. These results further link normative and neural decision-making while expanding our view of both as dynamic, adaptive processes that update and use expectations to govern both deliberation and commitment.

## Introduction

Even simple decisions can require us to adapt to a changing world. Should you go through the park orthrough town on yourwalk? The answer can depend on each route’s length, the weather, and/or the time of day. Some of these factors can change quickly and affect our deliberations in real time; e.g., an unexpected shower will send us hurrying down the faster route (***Figure 1A***), whereas spotting a new ice cream store can make the longer route more attractive. Despite the ubiquity of such dynamics in the real world, they are often neglected in models used to understand how the brain makes decisions. For example, many commonly used models assume that decision commitment occurs when the accumulated evidence for an option reaches a fixed, predefined value or threshold (*Wald, 1945; Ratcliff, 1978; Bogacz et al., 2006; Gold and Shadlen, 2007; Kilpatrick et al., 2019*). The value of this threshold can account for inherent trade-offs between decision speed and accuracy found in many tasks: lower thresholds generate faster, but less accurate decisions, whereas higher thresholds generate slower, but more accurate decisions (*Gold and Shadlen, 2007; Chittka et al., 2009; Bogacz et al., 2010*). However, these models do not adequately describe decisions made in environments with unknown or stochastically changing contexts (*Thura et al., 2014; Thura and Cisek, 2016; Palestro et al., 2018; Cisek et al., 2009; Drugowitsch et al., 2012; Thura et al., 2012; Tajima et al., 2019; Glickman et al., 2022*).

**Figure 1.**
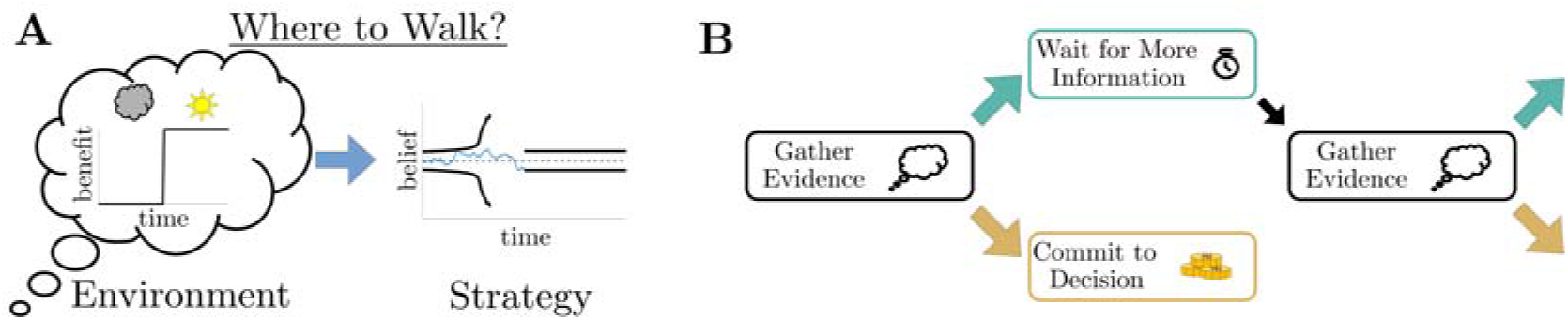
Simple decisions may require complex strategies. **A**: When choosing where to walk, environmental fluctuations (e.g., weather changes) may necessitate changes in decision bounds (black line) adapted to changes in the conditions (cloudy to sunny). **B**: Schematic of a dynamic programming. By assigning the best action to each moment in time, dynamic programming optimizes trial-averaged reward rate to produce the normative thresholds for a given decision.

Efforts to model decision-making thresholds under dynamic conditions have focused largely on heuristic strategies. For instance, “urgency-gating models” (UGMs) use thresholds that collapse monotonically over time (equivalent to dilating the belief in time) to explain decisions based on time-varying evidence quality (*Cisek et al., 2009; Carland et al., 2015; Evans et al., 2019*). Attempts to extend normative theory to such dynamic environments typically assume that individuals set decision thresholds to maximize trial-averaged reward rate (*Simen et al., 2009; Balci et al., 2011; Drugowitsch et al., 2012; Tajima et al., 2016; Malhotra et al., 2018; Boehm et al., 2020*), resulting in adaptive, time-varying thresholds similar to those assumed by heuristic UGMs. However, as in fixed-threshold models, these time-varying thresholds are typically defined before the evidence is accumulated, preceding the formative stages of the decision, and thus cannot account for environmental changes that may occur during deliberation.

To identify how environmental changes impact decision rules, we developed normative models of decision-making that adapt to dynamic changes in expectations or evidence. Specifically, we used Bellman’s equation (*Bellman, 1957; Mahadevan, 1996; Sutton et al., 1998; Bertsekas, 2012; Drugowitsch, 2015*) to identify decision strategies that maximize trial-averaged reward rate under dynamic conditions. We show that for simple tasks that include within-trial changes in the reward or the quality of observed evidence, these normative decision strategies involve non-trivial, time-dependent changes in decision thresholds. These rules take several different forms that outperform their heuristic counterparts, are identifiable from behavior, and have performance that is robust to noisy implementations. We also show that, compared to fixed-threshold models or UGMs, these normative, adaptive thresholds provide a better account of human behavior on a “tokens task,” in which both the value of commitment and evidence quality change at predictable times within each trial (*Cisek et al., 2009; Thura et al., 2014*). These results provide new insights into the behavioral relevance of a diverse set of adaptive decision thresholds in dynamic environments and tightly link the details of such environmental changes to threshold adaptations.

## Results

### Normative Theory for Dynamic Context 2AFC Tasks

To determine potential deliberation and commitment strategies used by subjects, we begin by identifying normative decision rules for two-alternative forced choice (2AFC) tasks with dynamic contexts. Normative decision rules that maximize trial-averaged reward rate can be obtained by solving an optimization problem using dynamic programming (*Bellman, 1957; Sutton et al., 1998; Drugowitsch et al., 2012; Tajima et al., 2016*). We define this trial-averaged reward rate, *ρ*, as (*Gold and Shadlen, 2002; Drugowitsch et al., 2012*)

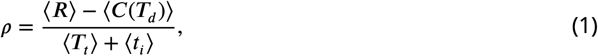

where 〈*R*〉 is the average reward for a decision, *T_d_* is the decision time, 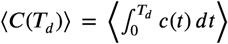 is the average total accumulated cost given an incremental cost function *c*(*t*), 〈***T**_t_*〉 is the average trial length, and (*t*) is the average inter-trial interval (*Drugowitsch, 2015*). Note that all averages in *Equation 1* are taken over trials. To find the normative decision thresholds that maximize *ρ*, we assign specific values (i.e., economic utilities) to correct and incorrect choices (reward and/or punishment) and the time required to arrive at each choice (i.e., evidence cost).

To represent the structure of a 2AFC tasks, we assume a decision environment for an observer with an initially unknown environmental state, *s* ∈ {*s*_+_, *s*_-_}, that uniquely determines which of two alternatives is correct. To infer the environmental state, this observer makes measurements, *ξ*, that follow a distribution *f*_±_(*ξ*) = *f* (*ξ*|*s*_±_) that depends on the state. Determining the correct choice is thus equivalent to determining the generating distribution, *f*_±_. An ideal Bayesian observer uses the log-likelihood ratio (LLR), *y*, to track their “belief” over the correct choice (*Wald, 1945; Bogacz et al., 2006; Veliz-Cuba et al., 2016*). After *n* discrete observations *ξ*_1:*n*_, the discrete-time LLR *y_n_* is given by

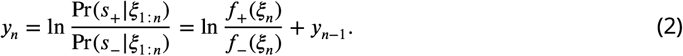

Given this defined task structure, we discretize the time during which the decision is formed and define the observer’s actions during each timestep. The observer gathers evidence (measurements) during each timestep prior to a decision and uses each increment of evidence to update their belief about the correct choice. Then, the observer has the option to either commit to a choice or make another measurement at the next timestep. By assigning a utility to each of these actions (i.e., a value *V*_+_ for choosing *s*_+_, a value *V*_-_ for choosing *s*_-_, and a value *V_w_* for sampling again), we can construct the value function for the observer given their current belief:

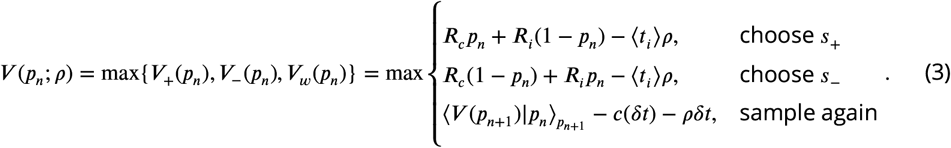

In *Equation 3*, 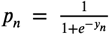 is the state likelihood at time *t_n_, **R**_c_* is the reward for a correct choice, *R_i_* is the reward for an incorrect choice, and *δt* is the timestep between observations. We choose generating distributions *f*_±_ that allow us to explicitly compute the average future value

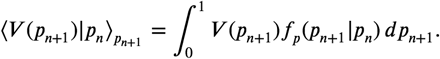

For a full derivation of *Equation 3*, as well as the detailed formulas necessary to compute this integral, see Methods and Materials. Using *Equation 3*, we find the specific belief values where the optimal action changes from gathering evidence to commitment, defining thresholds on the ideal observer’s belief that trigger decisions. *Figure 1***B** shows a schematic of this process.

To understand how normative decision thresholds adapt to fluctuating conditions, we derived them for several different forms of two-alternative forced-choice (2AFC) tasks in which we controlled changes in evidence or reward. For each task, the evidence was provided by observations drawn from a Gaussian distribution with one of two different means and signal-to-noise ratio (SNR) *m* (*Figure 2-Figure Supplement 1*). The SNR measures evidence quality: a smaller (larger) *m* implies that evidence is of lower (higher) quality, resulting in harder (easier) decisions. An observer must determine which of the two means were used to generate a finite number of observations. We introduced changes in the reward for a correct decision (“reward-change task”) or the SNR (“SNR-change task”) within a single decision, where the time and magnitude of the changes are known in advance to the observer (*Figure 1***A**, *Figure 2-Figure Supplement 2*). For example, changes in SNR arise naturally throughout a day as animals choose when to forage and hunt given variations in light levels and therefore target-acquisition difficulty (*Combes et al., 2012; Einfalt et al., 2012).*

Under these dynamic conditions, dynamic programming produces normative thresholds with rich non-monotonic dynamics (*Figure 2* **A,B**, *Figure 2-Figure Supplement 2*). For the reward-change task, these normative threshold dynamics exhibited several motifs that in some cases resembled fixed or collapsing thresholds characteristic of previous decision models, but in other cases exhibited novel dynamics. We characterized five different dynamic motifs in response to single changes in expected reward for different combinations of pre- and post-change reward values (*Figure 2***C** and **i-v**). For tasks in which reward is initially very low, thresholds are infinite until the reward increases, ensuring that the observer waits for the larger payout regardless of how strong their belief is (*Figure 2***i**). In contrast, when reward is initially very high, thresholds collapse to zero just before the reward decreases, ensuring all responses occur while payout is high (*Figure 2***v**). Between these two extremes, optimal thresholds exhibit rich, non-monotonic dynamics (*Figure 2***ii,iv**), promoting early decisions in the high-reward regime, or preventing early, inaccurate decisions in the low-reward regime. *Figure 2***C** shows the regions in pre- and post-change reward space where each motif is optimal, including broad regions with non-monotonic thresholds. Thus, even simple context dynamics can evoke complex decision strategies in ideal observers that differ from those predicted by constant decision-thresholds and heuristic UGMs.

**Figure 2.**
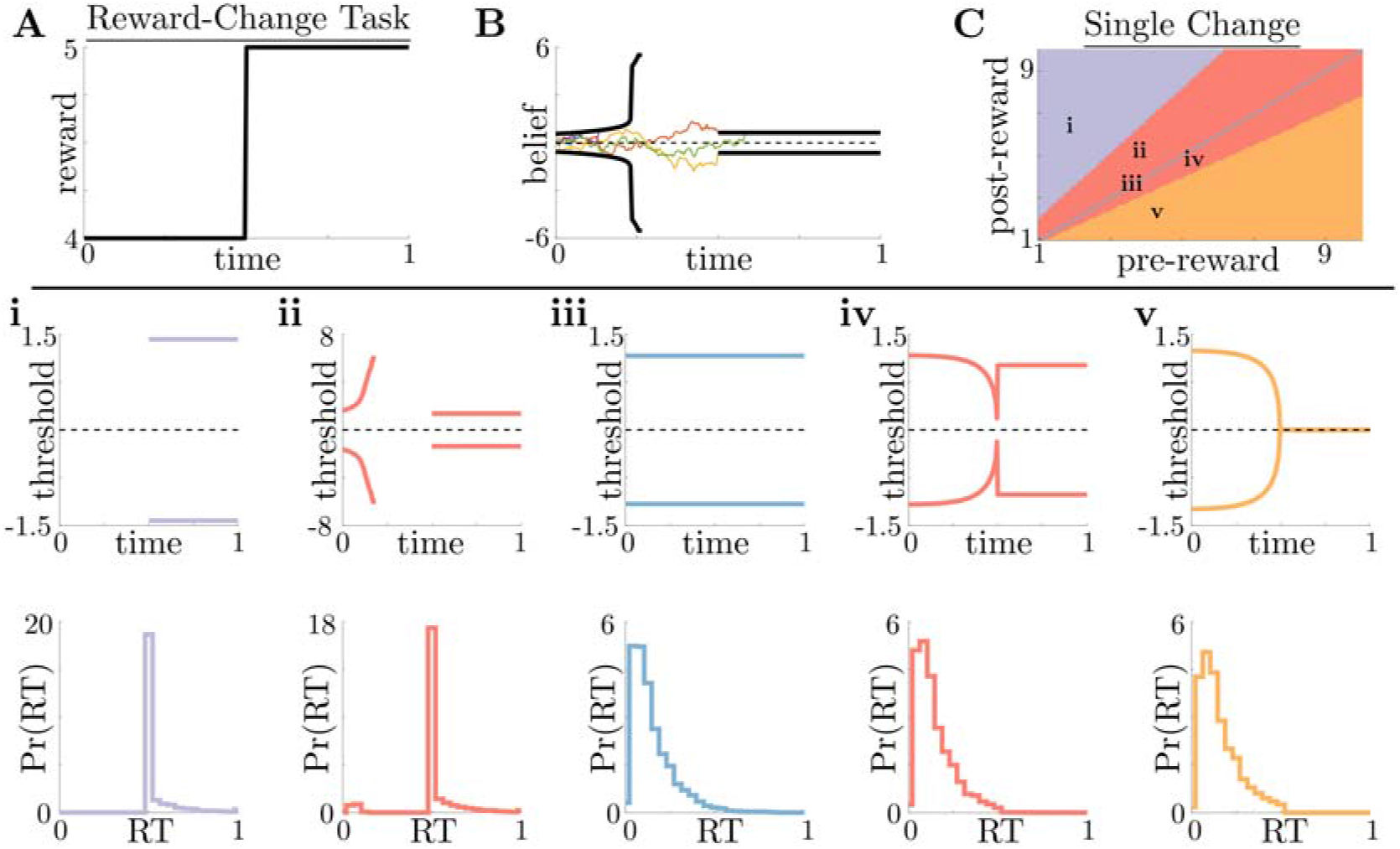
Normative decision rules are characterized by non-monotonic task-dependent motifs. **A,B**: Example reward time series for a reward-change task (black lines in **A**), with corresponding thresholds found by dynamic programming (black lines in **B**). The colored lines in **B** show sample realizations of the observer’s belief. **C**: To understand the diversity of threshold dynamics, we consider the simple case of a single change in the reward schedule. The panel shows a colormap of normative threshold dynamics for these conditions. Distinct threshold motifs are color-coded, corresponding to examples shown in panels **i-v. i-v**: Representative thresholds (top) and empirical response distributions (bottom) from each region in **C**. During times at which thresholds in the upper panels are not shown (e.g., *t* ∈ [0,0.5] in **i**), the thresholds are infinite and the observer will never respond. **Figure 2–Figure supplement 1.** Impact of evidence quality on belief and difficulty. **Figure 2–Figure supplement 2.** Normative thresholds for reward-change task with multiple changes. **Figure 2–Figure supplement 3.** Threshold dynamics in the inferred reward-change task.

We also formulated an “inferred reward-change task”, in which reward fluctuations are governed by a two-state Markov process and the observer infers these changes on-line. For this task, decision thresholds always changed monotonically with monotonic shifts in expected reward (see *Figure 2-Figure Supplement 3*). These results contrast with our findings with the reward-change task in which changes can be anticipated and monotonic changes in reward can produce nonmonotonic changes in decision thresholds.

For the SNR-change task, optimal strategies are characterized by threshold dynamics adapted to changes in evidence quality in a way similar to changes in reward (*Figure 3***A,B**, *Figure 3-Figure Supplement 1*). However, in this case monotonic changes in evidence quality always produce monotonic changes in response behavior. This observation holds across all of parameter space for evidence-quality schedules with single change points (**Figure 3C**), with only three optimal behavioral motifs (*Figure 3***i-iii**). This contrasts with our findings in the reward-change task, where monotonic changes in reward can produce non-monotonic changes in decision thresholds. Strategies arising from known dynamical changes in context tend to produce sharper response distributions around reward changes than around quality changes, which may be measurable in psychophysical studies. These findings suggest that changes in reward can have a larger impact on the normative strategy thresholds than changes in evidence quality.

**Figure 3.**
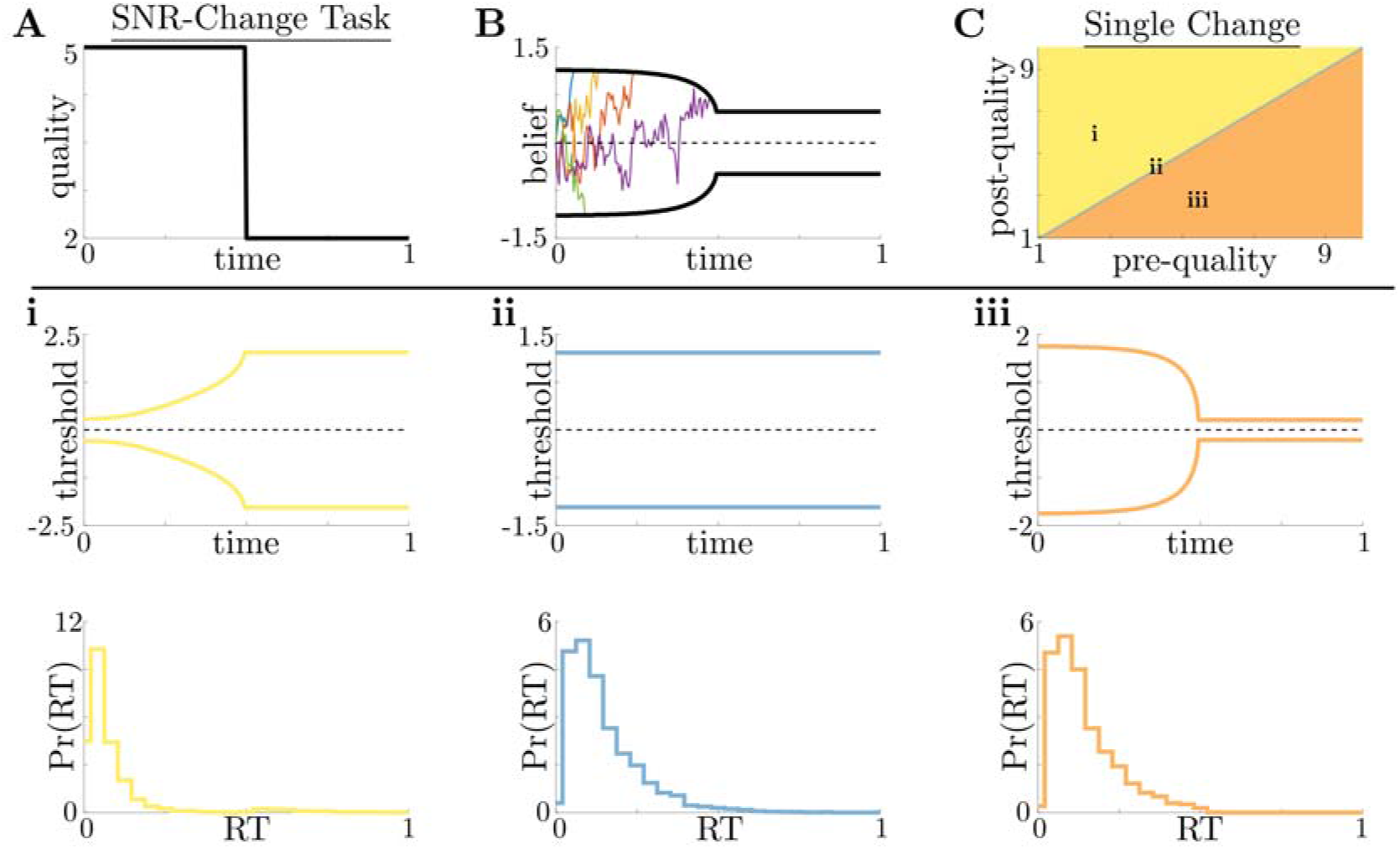
Dynamic-quality task does not exhibit non-monotonic motifs. **A,B**: Example quality time series for the SNR-change task (**A**), with corresponding thresholds found by dynamic programming (**B**). Colored lines in **B** show sample realizations of the observer’s belief. As in *Figure 2*, we characterize motifs in the threshold dynamics and response distributions based on single changes in SNR. **C**: Colormap of normative threshold dynamics for a known reward schedule task with a single quality change. Distinct dynamics are color-coded, corresponding to examples shown in panels **i**-**iii. i-iii**: Representative thresholds (top) and empirical response distributions (bottom) from each region in **C**. **Figure 3-Figure supplement 1.** Normative thresholds for SNR-change task with multiple changes.

#### Performance and Robustness of Non-monotonic Normative Thresholds

The normative solutions that we derived for dynamic-context tasks by definition maximize reward rate. This maximization assumes that the normative solutions are implemented perfectly. However, a perfect implementation may not be possible, given the complexity of the underlying computations, biological constraints on computation time and energy (*Louie et al., 2015*), and the synaptic and neural variability of cortical circuits (*Ma and Jazayeri, 2014; Faisal et al., 2008*). Given these constraints, subjects may employ heuristic strategies like the UGM over the normative model if noisy or mistuned versions of both models result in similar reward rates. We used synthetic data to better understand the relative benefits of different imperfectly implemented strategies. Specifically, we corrupted the internal belief state and simulated response times with additive Gaussian noise (See *Figure 4-Figure Supplement 1***C**) for three models:

1. The continuous-time normative model with time-varying thresholds ±*θ*(*t*) from *Equation 3* and belief that evolves according to the stochastic differential equation

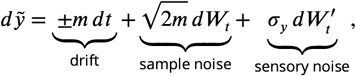

where *dW_t_* is a standard increment of a Wiener process, the sign of the drift ±*m dt* is given by the correct choice *s*_±_, and 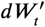 is an independent Wiener process with strength *σ_y_*. The addition of the additional noise process 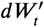 makes this a noisy Bayesian (NB) model.
2. A constant-threshold (Const) model, which uses the same belief 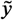 as the normative model but a constant, non-adaptive decision threshold ±*θ*(*t*) = ±*θ*_0_ (*Figure 4-Figure Supplement 1***A**).
3. The UGM, which uses the output of a low-pass filter as the belief,

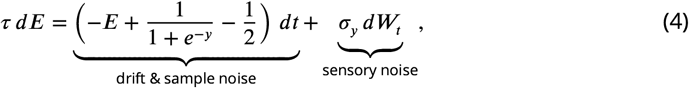

and commits to a decision when this output crosses a hyperbolically collapsing threshold 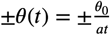 (*Figure 4-Figure Supplement 1***B**). In *Equation 4, E* is the filter’s output that serves as the UGM’s belief, *τ* is a relaxation time constant, and the noise-free observer’s belief *y* is the filter’s input.

For more details about these three models, see Methods and Materials. We compared their performance in terms of reward rate achieved on the same set of reward-change tasks shown in *Figure 2.*

When all three models were implemented without additional noise, the relative benefits of the normative model depended on the exact task condition. The performance differential between models was highest when reward changed from low to high values (*Figure 4***A**, dotted line; *Figure 4***B**). Under these conditions, normative thresholds are initially infinite and become finite after the reward increases, ensuring that most responses occur immediately once the high reward becomes available (*Figure 4***D**). In contrast, response times generated by the constant-threshold and UGM models tend to not follow this pattern. For the constant-threshold model, many responses occur early, when the reward is low (*Figure 4***E**). For the UGM, a substantial fraction of responses are late, leading to higher time costs (*Figure 4***F**). In contrast, when the reward changes from high to low values, all models exhibit similar response distributions and reward rates (*Figure 4***A**, dashed line; *Figure 4-Figure Supplement 2*). This result is not surprising, given that the constant-threshold model produces early peaks in the reaction time distribution, and the UGM was designed to mimic collapsing bounds that hasten decisions in response to imminent decreases in reward (*Cisek et al., 2009*). We therefore focused on the robustness of each strategy when corrupted by noise and responding to low-to-high reward switches -the regime differentiating strategy performance in ways that could be identified in subject behavior.

**Figure 4.**
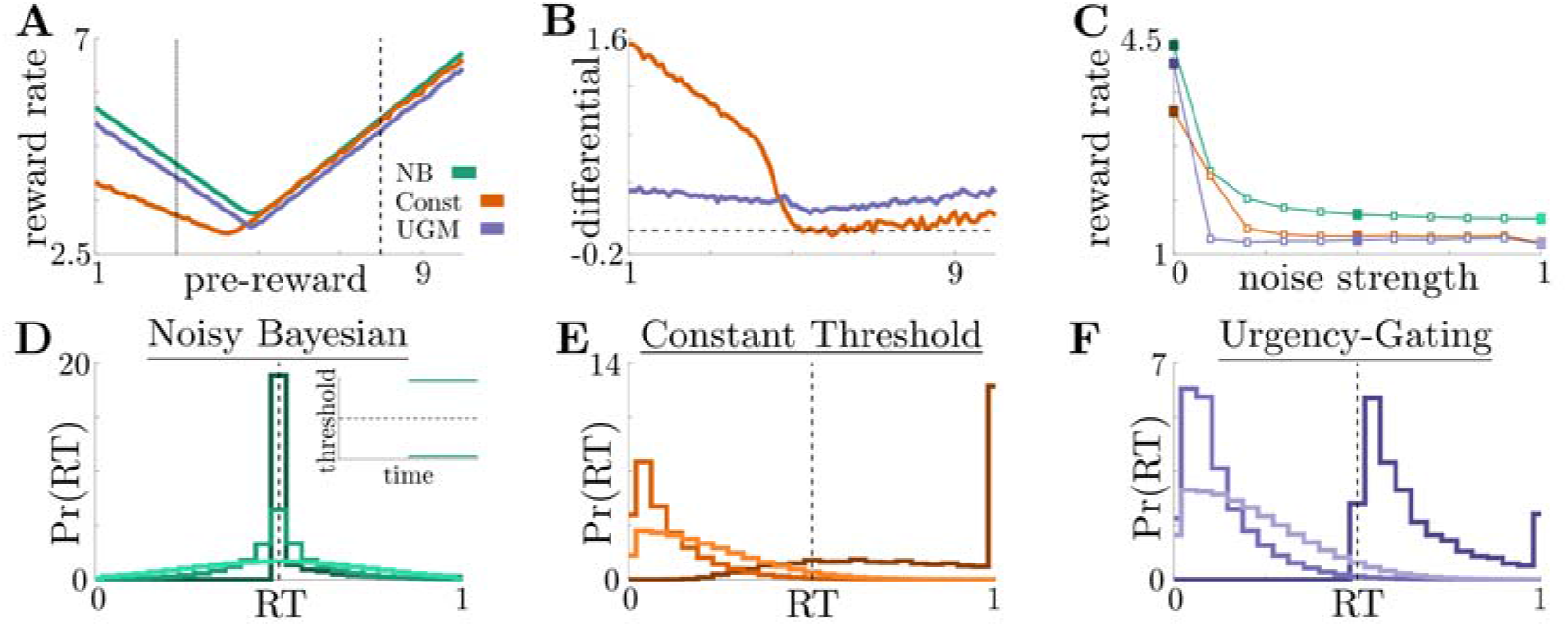
Benefits of adaptive normative thresholds compared to heuristics. **A**: Reward rate for the noisy Bayesian (NB) model, constant-threshold (Const) model, and UGM for the reward-change task given different pre-change rewards, with post-change reward set to keep the average total reward fixed (see Methods and Materials for details). Low-to-high reward changes (dotted line) produce larger performance differentials than high-to-low changes (dashed line). **B**: Absolute reward rate differential between NB and alternative models for different pre-change rewards. **C**: Reward rates of all models for low-to-high reward changes as both observation and response-time noise is increased (See *Figure 4-Figure Supplement 1***C** for reference) Filled markers correspond to no noise, moderate noise, and high noise strengths. **D,E,F**: Response distributions for (**D**) NB; (**E**) Const; and (**F**) UGM models in a low-to-high reward environment. In each panel, results derived for several noise strengths, corresponding with filled markers in **C**, are superimposed, with lighter distributions denoting higher noise. Inset in **D** shows normative thresholds obtained from dynamic programming. Dashed line shows time of reward increase. **Figure 4-Figure supplement 1.** Heuristic model and noise schematics. **Figure 4-Figure supplement 2.** Model performance for high-to-low reward switch. **Figure 4-Figure supplement 3.** Model performance for decomposed noise strengths.

Adding noise to the internal belief state (which tends to trigger earlier responses) and simulated response distributions (which tends to smooth out the distributions) does not alter the advantage of the normative model: across a range of added noise strengths, the normative model outperforms the other two when encountering low-to-high reward switches (*Figure 4***C**). This robustness arises because, prior to the reward change, the normative model uses infinite decision thresholds that prevent early noise-triggered responses when reward is low (*Figure 4***D**). In contrast, the heuristic models have finite collapsing or constant thresholds and thus produce more suboptimal early responses as belief noise is increased (*Figure 4***E,F**). Thus, adaptive decision strategies can result in considerably higher reward rates than heuristic alternatives even when implemented imperfectly, suggesting subjects may be motivated to learn such strategies.

### Adaptive Normative Strategies in the Tokens Task

To determine the relevance of the normative model to human decision-making, we analyzed previously collected data from a “tokens task” (*Cisek et al., 2009*). For this task, human subjects were shown 15 tokens inside a center target flanked by two empty targets (see *Figure 5***A** fora schematic). Every 200 ms, a token moved from the center target to one of the neighboring targets with equal probability. Subjects were tasked with predicting which flanking target would contain more tokens by the time all 15 moved from the center. Subjects could respond at any time before all 15 tokens had moved. Once the subject made the prediction, the remaining tokens would finish their movements to indicate the correct alternative. Given this task structure, one can show using a combinatorial argument (*Cisek et al., 2009*) that the state likelihood function *p_n_*, the probability the upper target will hold more tokens at the end of the trial, is given by

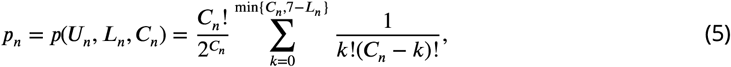

where *U_n’_ L_n’_* and *C_n_* are the number of tokens in the upper, lower, and center targets after token movement *n*, respectively. Because the total number of tokens was finite and known to the subject, token movements varied in their informativeness within a trial, yielding a dynamic and history-dependent evidence quality that, in principle, could benefit from adaptive decision processes (e.g., a token’s movement into a target is informative only if the difference in token counts between targets is lower than the number of tokens still in the center). In addition, the task included two different post-decision token movement speeds, “slow” and “fast”, that dynamically modulated the utility of decision commitment by altering the duration of the inter-trial interval, and hence the average rate at which rewards could be obtained. Given that costs and rewards can be subjective, we quantified how normative decision thresholds change with different combinations of rewards and costs, for both the slow (*Figure 5***B**) and fast (*Figure 5***C**) versions of the task.

**Figure 5.**
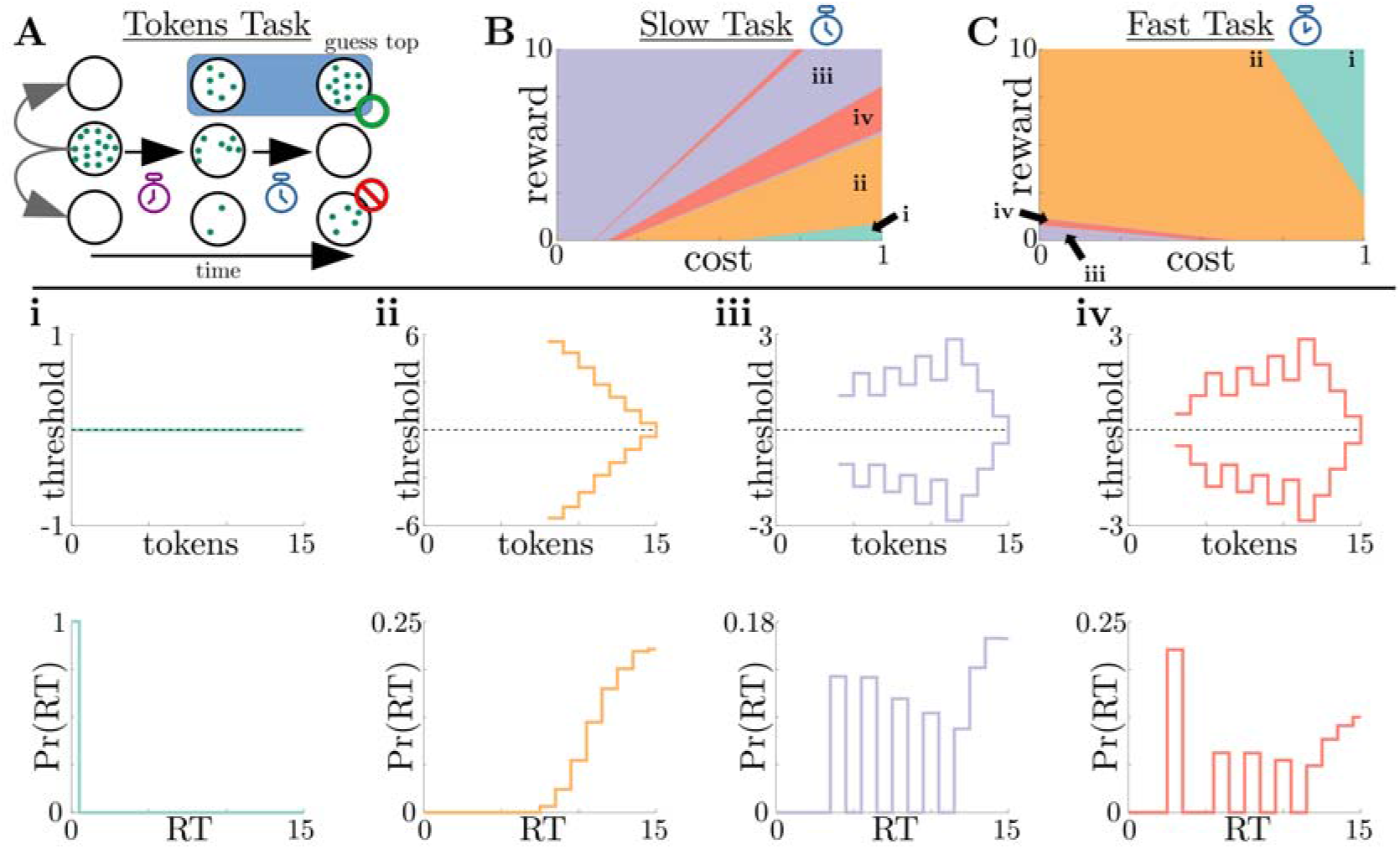
Normative strategies for the tokens task exhibit various distinct decision threshold motifs with sharp, non-monotonic changes. **A**: Schematic of the tokens task. The subject must predict which target (top or bottom) will have the most tokens once all tokens have left the center target (see text for details). **B**: Colormap of normative threshold dynamics for the “slow” version of the tokens task in reward-evidence cost parameter space (punishment *R_i_* is set to −1). Distinct dynamics are color-coded, with different motifs shown in **i-iv**. **C**: Same as **B**, but for the “fast” version of the tokens task. **i-iv**: Representative thresholds (top) and empirical response distributions (bottom) from each region in **B,C**. In regions where thresholds are not displayed (e.g., *N_t_* ∈ {0,...,7} in **ii**), the thresholds are infinite. **Figure 5-Figure supplement 1.** Tokens task thresholds in token lead space.

We identified four distinct motifs of normative decision threshold dynamics for the tokens task (*Figure 5***i-iv**). Some combinations of rewards and costs produced collapsing thresholds (*Figure 5***ii**) similar to the UGM developed by *Cisek et al. (2009)* for this task. In contrast, large regions of task parameter space produced rich non-monotonic threshold dynamics (*Figure 5***iii,iv**) that differed from any found in the UGM. In particular, as in the case of reward-change tasks, normative thresholds were often infinite for the first several token movements, preventing early and weakly informed responses. These motifs are similar to those produced by low-to-high reward switches in the reward-change task, but here resulting from the low relative cost of early observations. These non-monotonic dynamics also appear if we measure belief in terms of the difference in tokens between the top and bottom target, which we call “token lead space” (see *Figure 5-Figure Supplement 1*).

#### Adaptive Normative Strategies Best Fit Subject Response Data

To determine the relevance of these adaptive decision strategies to human behavior, we fit discrete-time versions of the noisy Bayesian (four free parameters), constant-threshold (three free parameters), and urgency-gating (five free parameters) models to response-time data from the tokens task collected by *Cisek et al. (2009)*. All models included belief and motor noise, as in our analysis of the dynamic-context tasks (*Figure 4-Figure Supplement 1***C**). The normative model tended to fit the data better than the heuristic models (see *Figure 6-Figure Supplement 1*), based on three primary analyses. First, both corrected AIC (AICc), which accounts for goodness-of-fit and model degrees-of-freedom, and average root-mean-squared error (RMSE) between the predicted and actual trial-by-trial response times, favored the noisy Bayesian model for most subjects for both the slow (*Figure 6***A**) and fast (*Figure 6***D**) versions of the task. Second, when considering only the bestfitting model for each subject and task condition, the noisy Bayesian model tended to better predict subject’s response times (*Figure 6***B,E**). Third, most subjects whose data were best described by the noisy Bayesian model had best-fit parameters that corresponded to non-monotonic decision thresholds, which cannot be produced by either of the other two models (*Figure 6***C,F**). Together, our results strongly suggest that these human subjects tended to use an adaptive, normative strategy instead of the kinds of heuristic strategies often used to model response data from dynamic context tasks.

**Figure 6.**
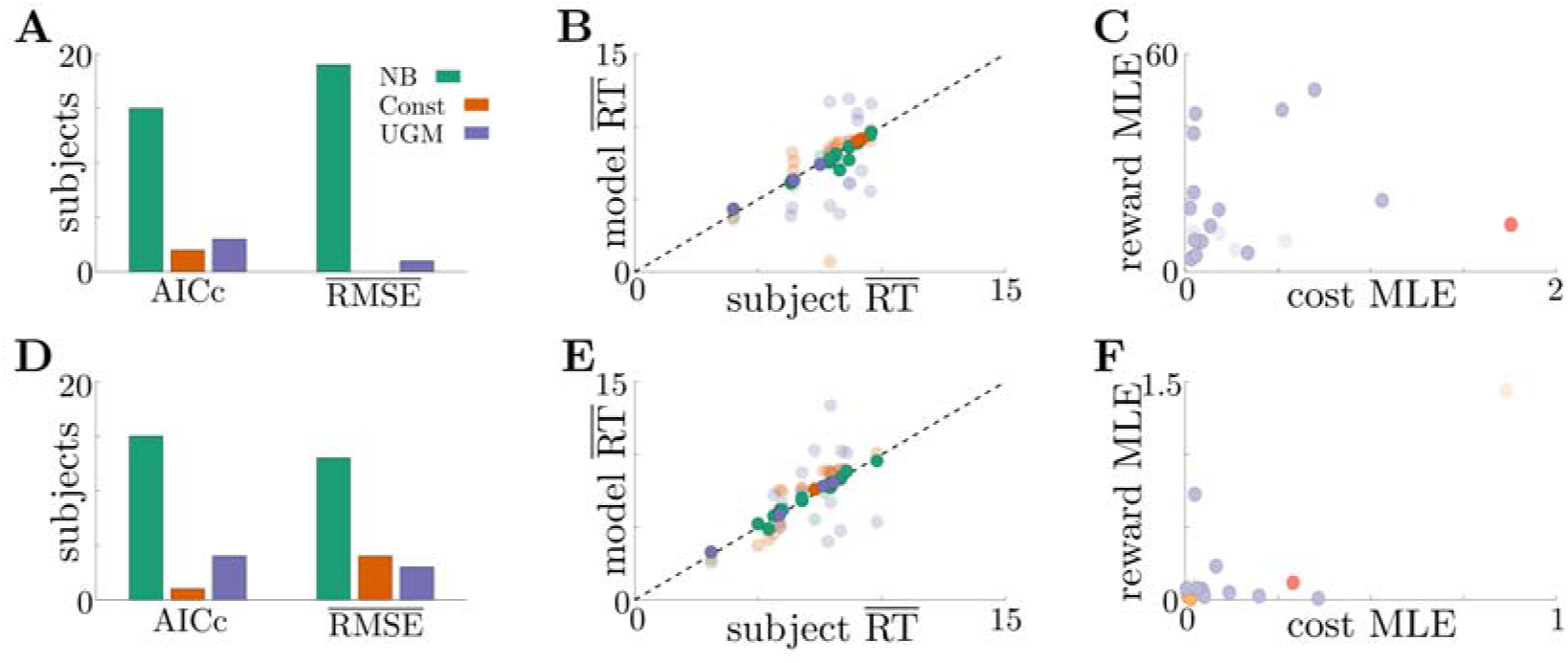
Adaptive normative strategies provide the best fit to subject behavior in the tokens task. **A**: Number of subjects from the slow version of the tokens task whose reponses were best described by each model (legend) identified using corrected AIC (left) and average trial-by-trial RMSE (right). **B**: Comparison of mean RT from subject data in the slow version of the tokens task (*x*-axis) to mean RT of each fit model (*y*-axis) at maximum-likelihood parameters. Each symbol is color-coded to agree with its associated model. Darker symbols correspond to the model that best describes the responses of a subject selected using corrected AIC. The NB model had the lowest variance in the difference between predicted and measured mean RT (NB var: 0.13, Const var: 3.11, UGM var: 5.39). **C**: Scatter plot of maximum-likelihood parameters for the noisy Bayesian model for each subject in the slow version of the task. Each symbol is color-coded to match the threshold dynamics heatmap from *Figure 5***B**. Darker symbols correspond to subjects whose responses were best described by the noisy Bayesian model using corrected AIC. **D-F**: Same as **A-C**, but for the fast version of the tokens task. The NB model had the lowest variance in the difference between predicted and measured mean RT in this version of the task (NB var: 0.22, Const var: 0.82, UGM var: 5.32). **Figure 6-Figure supplement 1.** Summary of model fits.

## Discussion

The goal of this study was to build on previous work showing that in dynamic environments, the most effective decision processes do not necessarily use relatively simple, pre-defined computations as in many decision models (*Bogacz et al., 2006; Cisek et al., 2009; Drugowitsch et al., 2012),* but instead adapt to learned or predicted features of the environmental dynamics (*Drugowitsch et al., 2014a*). Specifically, we used new “dynamic context” task structures to demonstrate that normative decision commitment rules (i.e., decision thresholds, or bounds, in “accumulate-to-bound” models) adapt to reward and evidence-quality switches in complex, but predictable, ways. Comparing the performance of these normative decision strategies to the performance of classic heuristic models, we found that the advantage of normative models is maintained when computations are noisy. We extended these modeling results to include the “tokens task”, in which evidence quality changes in a way that depends on stimulus history and the utility of commitment increases over time. We found that the normative decision thresholds for the tokens task are also non-monotonic and robust to noise. By reanalyzing human subject data from this task, we found most subjects’ response times were best-explained by a noisy normative model with non-monotonic decision thresholds. Taken collectively, these results show that ideal observers and human subjects use adaptive and robust normative decision strategies in relatively simple decision environments.

Our results can aid experimentalists investigating the nuances of complex decision-making in several ways. First, we demonstrated that normative behavior varies substantially across task parameters for relatively simple tasks. For example, the reward-change task structure produces five distinct behavioral motifs, such as waiting until reward increases (*Figure 2***i**) and responding before reward decreases unless the accumulated evidence is ambiguous (*Figure 2***iv**). Using these kinds of modeling results to inform experimental design can help us understand the possible behaviors to expect in subject data. Furthermore, extending our work and considering the sensitivity of performance to both model choice and task parameters (*Barendregt et al., 2019; Radillo et al., 2019*) will help to identify regions of task parameter space where models are most identifiable from observables like response time and choice. In general, our work suggests that experimentalists can design more informative tasks by using normative theory to determine what subject strategies are plausible, the volume and diversity of tasks needed to identify them, and the relationship between task dynamics and decision rules.

Real subjects likely do not rely on a single strategy when performing a sequence of trials (*Ashwood et al., 2022*) and instead rely on a mix of near-normative, sub-normative, and heuristic strategies. In fitting subject data, experimentalists are thus presented with the difficult task of constructing a library of possible models to use in their analysis. More general approaches have been developed for fitting response data to a broad class of models (*Shinn et al., 2020),* but these model libraries are typically built on pre-existing assumptions of how subjects accumulate evidence and make decisions. Because the potential library of decision strategies is theoretically limitless, a normative analyses can both expand and provide insights into the range of possible subject behaviors in a systematic and principled way. Understanding this scope will assist in developing a well-groomed candidate list of near-normative and heuristic models. For example, if a normative analysis of performance on a dynamic reward task produces threshold dynamics similar to those in *Figure 2***B**, then the fitting library should include a piecewise-constant threshold (or urgency signal) model. Combining these model-based investigations with model-free approaches, such as rate-distortion theory (*Berger, 2003; Eissa et al., 2021),* can also aid in identifying commonalities in performance and resource usage within and across model classes without the need for pilot experiments.

Our work complements the existing literature on optimal decision thresholds by demonstrating the prevalence of behaviors reflective of non-monotonic decision thresholds. Most studies describing decision strategies with time-varying decision thresholds focus on environments with fixed structure, in which dynamic decision thresholds are adapted as the observer acquires knowledge of the environment. Using dynamic programming (*Drugowitsch et al., 2012,2014b; Tajima et al., 2016*) or policy iteration (*Malhotra et al., 2017, 2018),* normative strategies in these environments typically have monotonically collapsing decision thresholds that can be approximated by a standard UGM (*Tajima et al., 2019).* While recent work has started to generalize notions of urgency-gating behavior (*Trueblood et al., 2021),* we have shown that novel response behaviors need to be considered even with simple tasks.

The neural mechanisms responsible for implementing and controlling decision thresholds are not well understood. Recent work has identified several cortical regions that may contribute to threshold formation, such as prefrontal cortex (*Hanks et al., 2015),* dorsal premotor area (*Thura and Cisek, 2020*), and superior colliculus (*Crapse et al., 2018; Jun et al., 2021).* Urgency signals are a complementary way of dynamically changing decision thresholds via a commensurate scale in belief, which *Thura and Cisek (2017)* suggest are detectable in recordings from basal ganglia. The normative decision thresholds we derived do not employ urgency signals, but analogous UGMs may involve non-monotonic signals. For example, the switch from an infinite-to-constant decision threshold typical of low-to-high reward switches would correspond to a signal that suppresses responses until a reward change. Measurable signals predicted by our normative models would therefore correspond to zero mean activity during low reward, followed by constant mean activity during high reward. While more experimental work is needed to test this hypothesis, our work has expanded the view of normative and neural decision making as dynamic processes for both deliberation and commitment.

## Methods and Materials

### Normative Decision Thresholds from Dynamic Programming

Here we detail the dynamic programming tools required to find normative decision thresholds. For the free-response tasks we consider, an observer sets their potentially time-dependent decision thresholds, *θ*_±_(*t*), that determine when they will stop accumulating evidence and commit to a choice: When *y* ≥ *θ*_+_(*t*) (*y* ≤ *θ*_-_(*t*)), the observer chooses the state *s*_+_ (*s*_-_). In general, an observer is free to set *θ*_±_(*t*) any way they wish. However, a normative observer sets these thresholds to optimize an objective function, which we assume throughout this study to be the trial-averaged reward rate, *p*, which is given by *Equation 1.* In this definition of reward rate, the incremental cost function *c(t)* accounts for both explicit costs (e.g., paying for observed evidence, metabolic costs of storing belief in working memory) and implicit costs (e.g., opportunity cost). We assume symmetry in the problem (in terms of prior, rewards, etc.) that guarantees the thresholds are symmetric about *y* = 0 and *θ*_±_(*t*) = ±*θ*(*t*). We derive the optimal threshold policy for a general incremental cost function *c*(*t*), but in our results we consider only constant costs functions *c*. Although the space of possible cost functions is large, restricting to a constant value ensures that threshold dynamics are governed purely by task and reward structure and not by an arbitrary evidence cost function.

To find the thresholds ±*θ* that optimize the reward rate given by *Equation 1*, we start with a discrete-time task where observations every *δt* time units, and we simplify the problem so the length of each trial is fixed and independent of the decision time *T_d_*. This simplification makes the denominatorof *ρ* constantwith respect to trial-to-trial variability, meaning we can optimize reward rate by maximizing the numerator 〈*R*〉 – 〉*C*(*T_d_*)〉. Under this simplified task structure, we suppose the observer has just drawn a sample *ξ_n_* and updated their state likelihood to 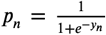. At this moment, the observer takes one of three possible actions:

1. *Stop accumulating evidence and commit to choice s*_+_. This action has value equal to the average reward for choosing *s*_+_, which is given by

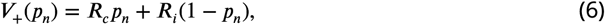

where *R_c_* is the value for a correct choice and *R_t_* is the value for an incorrect choice.
2. *Stop accumulating evidence and commit to choice s_-_*. By assuming the reward for correctly (or incorrectly) choosing *s*_+_ is the same as choosing *s*_-_, the value of this action is obtained by symmetry from *Equation 6*:

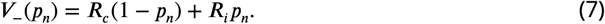
3. *Wait to commit to a choice and draw an additional piece of evidence.* Choosing this action means the observer expects their future overall value *V* to be greater than their current value, less the cost incurred by waiting for additional evidence. Therefore, the value of this choice is given by

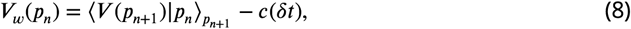

where *c* is the incremental evidence cost function; because we assume that the incremental cost is constant, this simplifies *c*(*δt*) = *cδt*.

Given the action values from *Equation 6-Equation 8*, the observer takes the action with maximal value, resulting in their overall value function

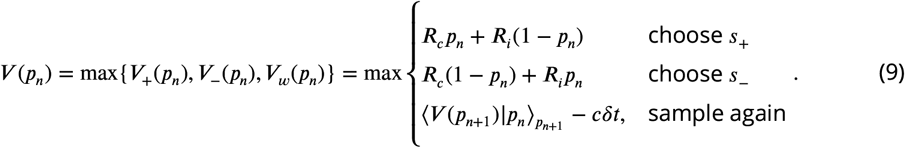

Because the value-maximizing action depends on the state likelihood, *p_n_*, the regions of likelihood space where each action is optimal divide the space into three disjoint regions. The boundaries of these regions are exactly the optimal decision thresholds, which can be mapped to LLR-space to obtain _±_*θ*. To find these thresholds numerically, we used backward induction starting at the total trial length *t* = *T_t_*. At this moment in time, it impossible to wait for more evidence, so the value function in *Equation 9* does not depend on the future. Once the value is calculated at this time point, it can be used as the future value at time point *t* = *T_t_* – *δt*.

To find the decision thresholds for the desired tasks where *T_t_* is not fixed, we must optimize both the numerator and denominator of *Equation 1.* To account for the variable trial length, we adopt techniques from average reward reinforcement learning (*Mahadevan, 1996)* and penalize the waiting time associated with each action by the waiting time itself scaled by the reward rate *ρ* (i.e., 〈*t_i_*〉_*ρ*_ for committing to *s*_+_ or *s*_-_ and *ρδt* for waiting). This modification makes all trials effectively the same length and allows us to use the same approach used to derive *Equation 9* (*Drugowitsch et al., 2012*). The new overall value function is given by *Equation 3*:

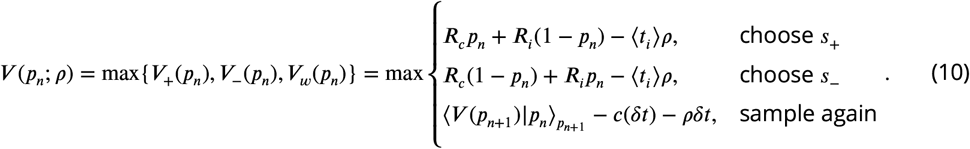

To use this new value function to numerically find the decision thresholds, we must note two new complications that arise from moving away from fixed-length trials. First, we no longer have a natural end time from which to start backward induction. We remedy this issue by following the approach of *Drugowitsch et al. (2012)* and artificially setting a final trial time *T_f_* that is far enough in the future so that decision times of this length are highly unlikely and do not impact the response distributions. If we desire accurate thresholds up to a time *t*, we set *T_f_* = 5*t*, which produces an accurate solution while avoiding a large numerical overhead incurred from a longer simulation time. In our simulations, we set *t* based on when we expect most decisions to be made. Second, the value function now depends on the unknown quantity *ρ*, resulting in a co-optimization problem. To address this complication, note that when *ρ* is maximized, our derivation requires *V*(0; *ρ*) = 0 for a consistent Bellman’s equation (*Drugowitsch et al., 2012*). We exploit this consistency requirement by fixing an initial reward rate *ρ*_0_, solving the value function through backward induction, calculating *V*(0; *ρ*_0_), and updating the value of *ρ* via a root finding scheme. For more details on numerical implementation, see https://github.com/nwbarendregt/AdaptNormThresh.

### Dynamic Context 2AFC Tasks

For all dynamic context tasks, we assume that observations follow a Gaussian distribution with so that 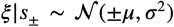. Using the Functional Central Limit Theorem, one can show (*Bogacz et al., 2006*) that in the continuous-time limit, the belief *y* evolves according to a stochastic differential equation:

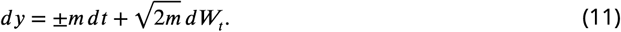

In *Equation 11*, 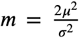 is the scaled signal-to-noise ratio (SNR), *dW_t_* is a standard increment of a Wiener process, and the sign of the drift ±*m dt* is given by the sign of the correct choice *s*_±_. To construct Bellman’s equation for this task, we start by discretizing time *t*_1:*n*_ and determine the average value gained we must also determine the average value gained by waiting and collecting another observation at *t*_*n*+1_:

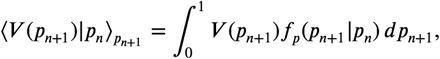

where *p_n_* is the probability the environment is in state *s*_+_. The main difficulty in computing this expectation is computing the likelihood transfer function *f_p_*(*p*_*n*+1_|*p_n_*). To compute this transfer function, we can start by using the definition of the LLR *y_n_* and leveraging the relationship between *p_n_* and *y_n_* to find *p*_n_ and a function of the observation *ξ_n_*:

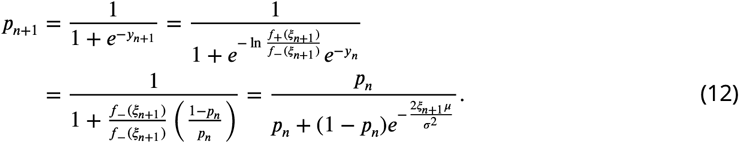

Note that we used the fact that in continuous-time, the observations 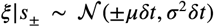. The relationship between *ξ*_*n*+1_ and *p*_*n*+1_ in Equation 12 can be inverted to obtain

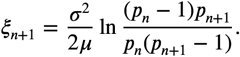

With this relationship established, we can find the likelihood transfer function 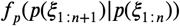 by finding the observation transfer function 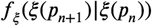 and performing a change of variables, which by independence of the sample is simply *f_ξ_*(*ξ*_*n*+1_). With probability *p_n_*, *ξ*_*n*+1_ will be drawn from the normal distribution 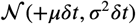, and with probability 1 – *p_n_*, *ξ*_*n*+1_ will be drawn from the normal distribution 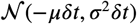. This immediately provides the observation transfer function by marginalizing:

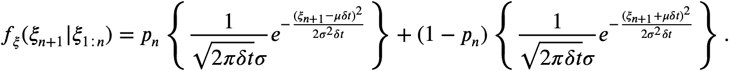

Performing the change of variables using the derivative 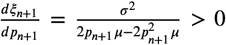 yields the transfer function

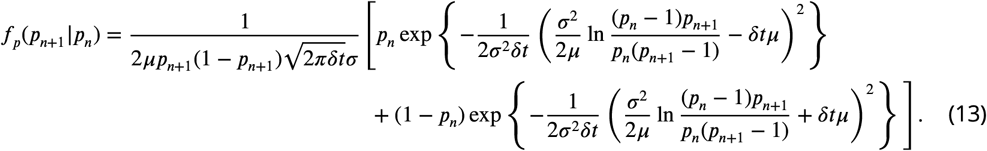

Combining *Equation 11* and *Equation 13*, we can construct Bellman’s equation for any dynamic context task.

#### Reward-Change Task Thresholds

For the reward-change task, we fixed punishment ***R**_i_*, = 0 and allowed the reward ***R**_c_* to be a Heavi side function:

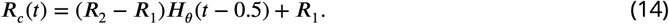

In *Equation 14*, there is a single switch in rewards between pre-change reward ***R***_1_ and post-change reward ***R***_2_. This change occurs at *t* = 0.5. Substituting this reward function into Equation 3 allows us to find the normative thresholds for this task as a function of *R*_1_ and *R*_2_.

For the inferred reward change task, we allowed the reward ***R***(*t*) ∈ {***R**_H_,**R**_L_*} to be controlled by a continuous-time two-state Markov process with transition (hazard) rate *h* between rewards ***R**_H_* ≥ ***R***_*L*_. In addition, the state of this Markov process must be inferred from an independent evidence source to the environment’s state (i.e., the correct choice); for simplicity, we assume that the reward-evidence source is also Gaussian-distributed with quality 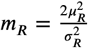. Glaze et al. (2015); *Veliz-Cuba et al. (2016); Barendregt et al. (2019)* have shown that the belief *y_R_* for such a dynamic state inference process is given by the modified DDM

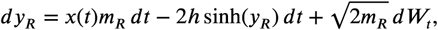

where *x*(*t*) ∈ ± 1 is a telegraph process that mirrors the state of the reward process (i.e., *x*(*t*) = 1 when ***R***(*t*) = ***R***_*H*_ and *x*(*t*) = – 1 when ***R***(*t*) = ***R***_*L*_). With this belief over reward state, we must also modify the values ***V***_+_(*p_n_*) and ***V***_-_(*p_n_*) to account for the uncertainty in ***R**_c_*. Defining 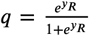 as the reward likelihood gives

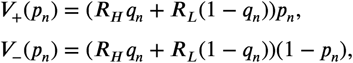

where we have fixed ***R**_i_* = 0 for simplicity.

#### SNR-Change Task Thresholds

For the SNR-change task, we allowed the task difficulty *m* to vary over a single trial by making *μ*(*t*) a time-dependent step function similar to *Equation 14*:

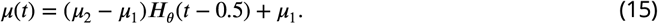

In *Equation 15*, there is a single switch in evidence quality between pre-change quality *μ*_1_ and postchange quality *μ*_2_. This change occurs at *t* = 0.5. Substituting this quality time series into the likelihood transfer function in *Equation 13* allows us to find the normative thresholds for this task as a function of *μ*_1_ and *μ*_2_. This modification necessitates that the transfer function *f_p_* also be a function of time; however, because the quality change points are known in advance to the observer, we can simply change between different transfer functions at the specified quality changes.

### Reward-Change Task Model Performance

Here we detail the three models used to compare observer performance in the reward-change task, as well as the noise filtering process used to generate synthetic data. For the noisy Bayesian model, the observer uses the thresholds ±*θ*(*t*) obtained via dynamic programming, thus making the observer a noisy ideal observer. For the constant-threshold model, the observer uses a constant threshold ±*θ*(*t*) = ±*θ*_0_, which is predicted to be optimal only in simple, static decision environments. Both the noisy Bayesian and constant-threshold models also use a noisy perturbation of the LLR 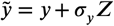 as their belief, where *σ_y_* is the strength of the noise and *Z* is a sample from a standard normal distribution. In continuous-time, this perturbation involves adding an independent Wiener process to *Equation 11*:

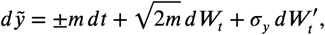

where 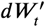 is an independent Wiener process with strength *σ_y_*. The UGM, being a phenomenological model, behaves differently from the other models. The UGM belief *E* is the output of the noisy low-pass filter given by *Equation 4.* To add noise to the UGM’s belief variable *E*, we simply allowed *σ_y_* > 0 in the low-pass filter in *Equation 4*.

In addition to the inference noise, we also filtered each process through a Gaussian responsetime filter with strength *σ_mn_*, so that if the model predicted a response time *T*, the measured response time 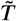 was drawn from a normal distribution centered at *T* with standard deviation *σ_mn_*.

To compare model performance on the reward-change task, we first fixed the value of pre-change reward ***R***_1_ (and set ***R***_1_ + ***R***_2_ = 11) to find the post-change reward) and tuned each model to achieve optimal reward rate with no additional noise in both the inference and response processes. Bellman’s equation outputs both the optimal normative thresholds and reward rate, allowing us to find the exact tuning of the normative model. For the constant threshold model and the UGM, we approximate optimal tuning by using a grid search over each models parameters. After tuning all models for a given reward structure, we filtered them through the two noise sources. When generating noisy synthetic data from these models, we generated 100 synthetic subjects, each with sampled noise strengths *σ_y_* and *σ_mn_*. We defined “noise strength” of noise samples (*σ_y’_*, *σ_mn_*) to be the ratio

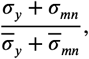

where 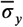 and 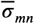 are the maximum values of belief noise and motor noise considered, respectively. Noise strength is thus defined between 0 and 1, such that a noise strength of 0.5 is approximately equivalent to the fitted noise strength obtained from tokens task subject data. We plot the response distributions using noise strengths of 0, 0.5, and 1 in our results. We then generated 1000 trials for each subject and had each simulated subject repeat the same block of trials three times, one for each model. This process ensured that the only difference between model performance would come from their distinct threshold behaviors, because each model was taken to be equally noisy and was run using the same stimuli.

### Tokens Task

#### Normative Model for the Tokens Task

For the tokens task, observations in the form of token movements are Bernoulli distributed with parameter *p* = 0.5 that occur every 200 ms. Because of the stimulus structure, one can show using a combinatorial argument (*Cisek et al., 2009)* that the likelihood function *p_n_* is given by *Equation 5.* Constructing the likelihood transfer function *f_p_* required for Bellman’s equation is also simplified from the Gaussian 2AFC tasks, as there are only two possible likelihoods that one can transition two after observing a token movement:

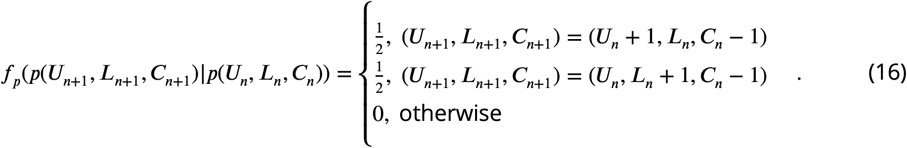

Combining *Equation 5* and *Equation 16*, we can fully construct Bellman’s equation for the tokens task. While the timings of the token movements, post-decision token acceleration, and inter-trial interval are fixed, we let the reward ***R**_c_* and cost function *c* be free parameters to control the different threshold dynamics of the model.

#### Model Fitting and Comparison

We used three models to fit the subject response data provided by *Cisek et al. (2009)*: the noisy Bayesian model (*k* = 4 parameters), the constant threshold model (*k* = 3 parameters), and the UGM (*k* = 5) parameters. To fit each model, we used Markov Chain Monte Carlo (MCMC) with a standard Gaussian proposal distribution to generate an approximate posterior made up of 10,000 samples. For more details as to our specific implementation of MCMC for this data, see the MATLAB code available at https://github.com/nwbarendregt/AdaptNormThresh. We held out 2 of the 22 subjects to use as training data when tuning the covariance matrix of the proposal distribution for each model, and performed the model fitting and comparison analysis on the remaining 20 subjects. Using the approximate posterior obtained via MCMC for each subject and model, we used calculated AICc using the formula

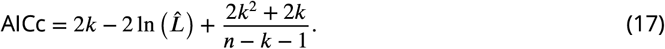

In *Equation 17, k* is the number of parameters of the model, 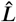 is the likelihood of the model evaluated at the maximum-likelihood parameters, and *n* is the number of responses in the subject data (*Cavanaugh, 1997; Brunham and Anderson, 2002*). Because each subject performed different numbers of trials, using AICc allowed us to normalize results to account for the different data sizes; note that for many responses (i.e., for large *n*), AICc converges to the standard definition of AIC. For the second model selection metric, we measured how well each fitted model predicted the trial-by-trial responses of the data by calculating the average RMSE between the response times from the data and the response times predicted by each model. To measure the difference between a subject’s response time distribution and the fitted model’s distribution (*Figure 6-Figure Supplement 1*), we used Kullback-Leibler (KL) divergence:

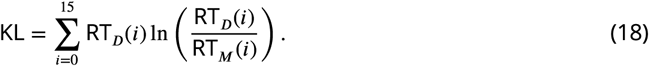

In *Equation 18, i* is a time index representing the number of observed token movements, RT_*D*_(*i*) is the probability of responding after *i* token movements from the subject data, and RT_*M*_(*i*) is the probability of responding after *i* token movements from the model’s response distribution. Smaller values of KL divergence indicate that the model’s response distribution is more similar to the subject data.

## Code Availability

See https://github.com/nwbarendregt/AdaptNormThresh for the MATLAB code used to generate all results and figures.

## Acknowledgments

We thank Paul Cisek for providing response data from the tokens task used in our analysis.

---

**Figure 2-Figure supplement 1.**
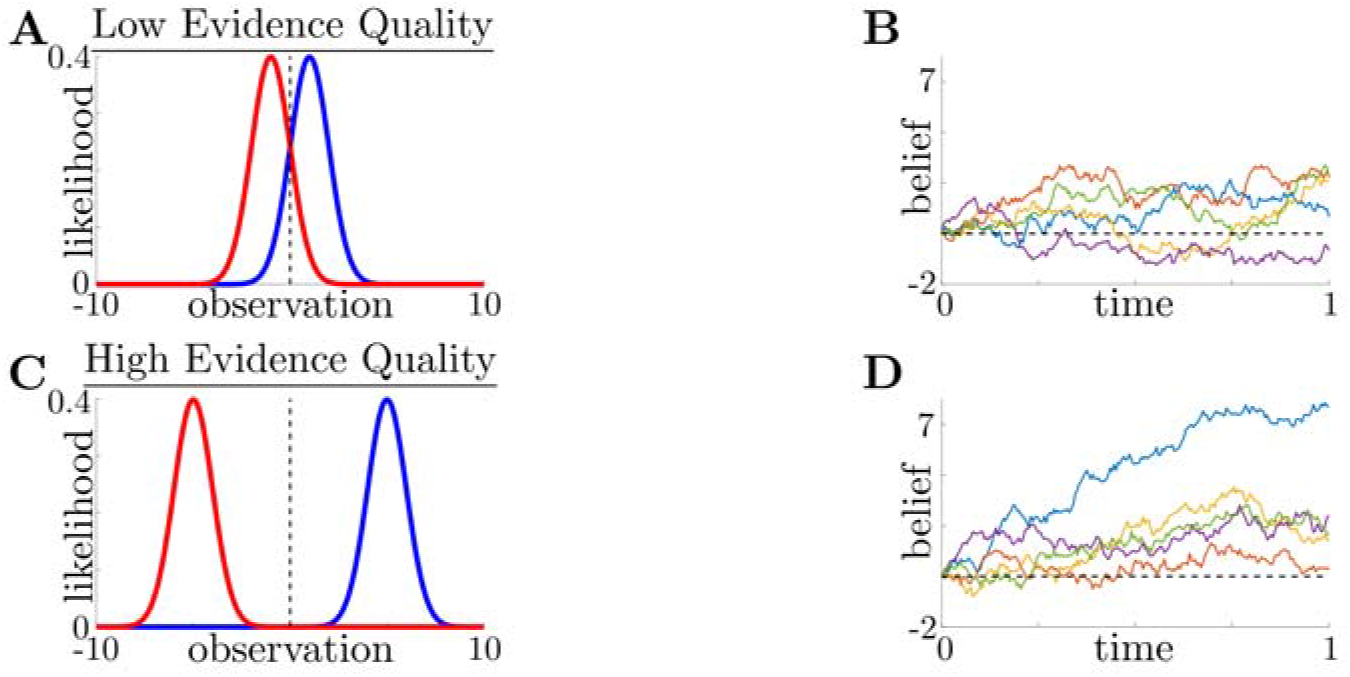
Evidence quality impacts observation distinguishability and task difficulty. **A:** Likelihood functions for environmental states (i.e., possible choices) *s*_+_ (blue) and *s*___ (red) in a low evidence quality task (*m* = 2), where we define 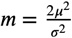, a scaled signal-to-noise ratio, as the evidence quality. **B**: Observer belief (LLR of accumulated evidence) in a low evidence quality task, with several belief realizations superimposed. **C,D**: Same as **A** and **B**, but for a high evidence quality task (*m* = 50).

**Figure 2-Figure supplement 2.**
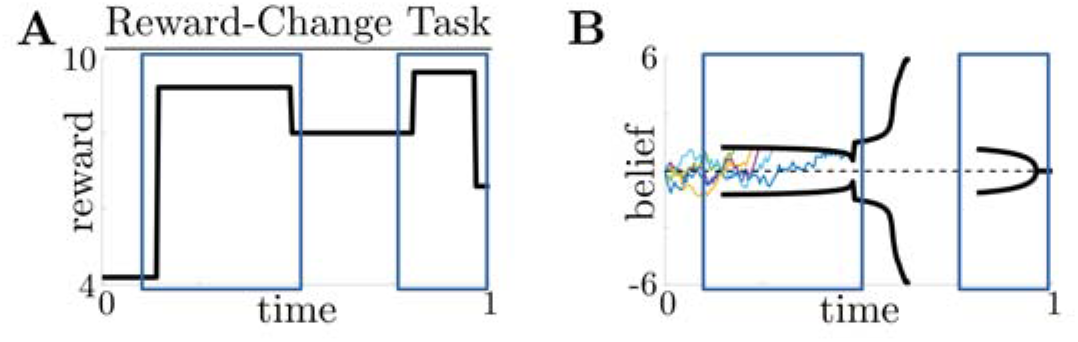
Normative decision thresholds exhibit multiple motifs for multiple reward changes. **A,B**: Example reward time series for a reward-change task (black lines in **A**), with corresponding thresholds found by dynamic programming (black lines in **B**). The colored lines in **B** show sample realizations of the observer’s belief. Similar changes in reward (boxed regions) produce similar motifs in threshold dynamics.

**Figure 2-Figure supplement 3.**
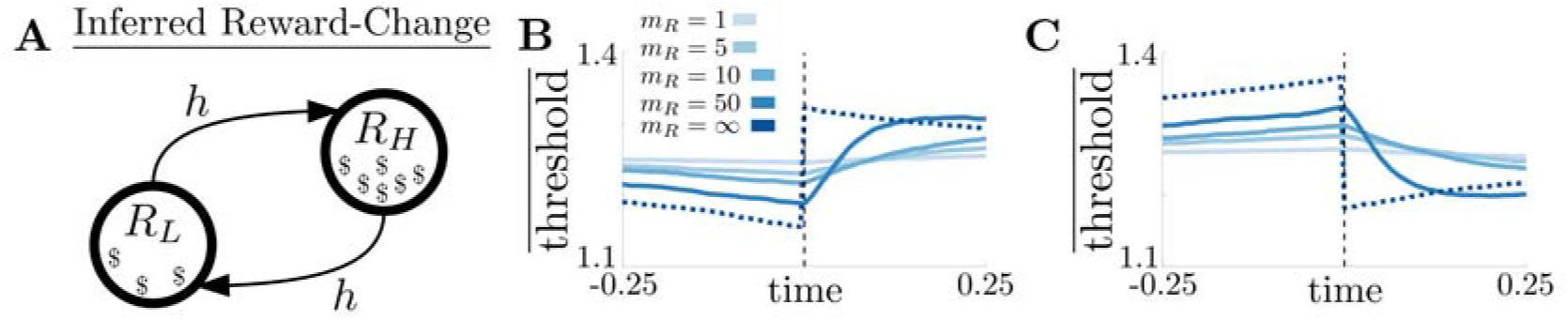
Threshold dynamics in the inferred reward change task track piecewise constant dynamics. **A**: Markov process governing reward states with rewards *R_L_* ≤ *R_H_* and symmetric transition (hazard) rate *h* between states. **B**: Change point-triggered average of normative thresholds for a high-to-low reward change. Several values of reward inference difficulty *m_R_* are superimposed (legend). Dotted line corresponds to the thresholds for an infinite *m_R_* task. **C**: Same as **B**, but for a low-to-high reward change.

**Figure 3-Figure supplement 1.**
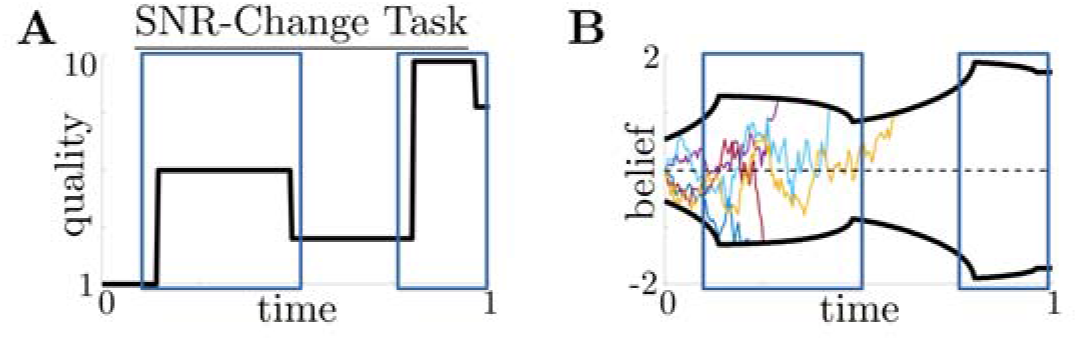
Normative decision thresholds change monotonically when anticipating SNR changes. **A,B**: Example quality time series for the SNR-change task with multiple changes (**A**), with corresponding thresholds found by dynamic programming (**B**). Colored lines in (**B**) show sample realizations of the observer’s belief. Similar changes in quality (boxed regions) produce similar motifs in threshold dynamics.

**Figure 4-Figure supplement 1.**
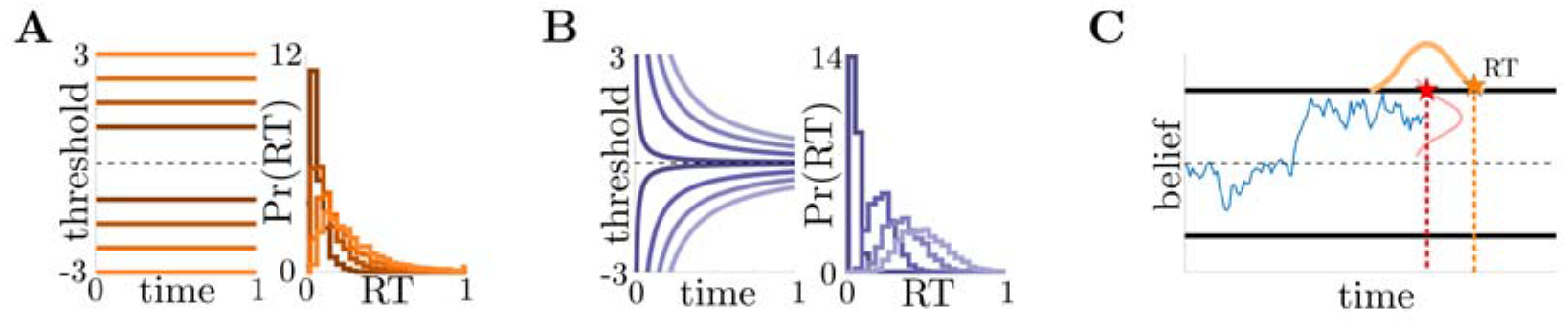
Heuristic observer modelsand noise filters. **A:** Constantthreshold (Const) model in belief-space (left), with associated RT distributions (right). Several versions of model are superimposed (gradient). **B**: Threshold dynamics of the UGM (left), with associated RT distributions (right). The UGM uses a low-pass filtered version of the normative LLR as its belief and dilates this belief linearly in time; this is equivalent to hyperbolically-collapsing decision thresholds (see Methods and Materials). **C**: Schematic of noise generation processes, describing possible sources of performance decreases. Gaussian noise with standard deviation *σ_y_* is added to belief (red) and with standard deviation *σ_mn_* added to response times (orange).

**Figure 4-Figure supplement 2.**
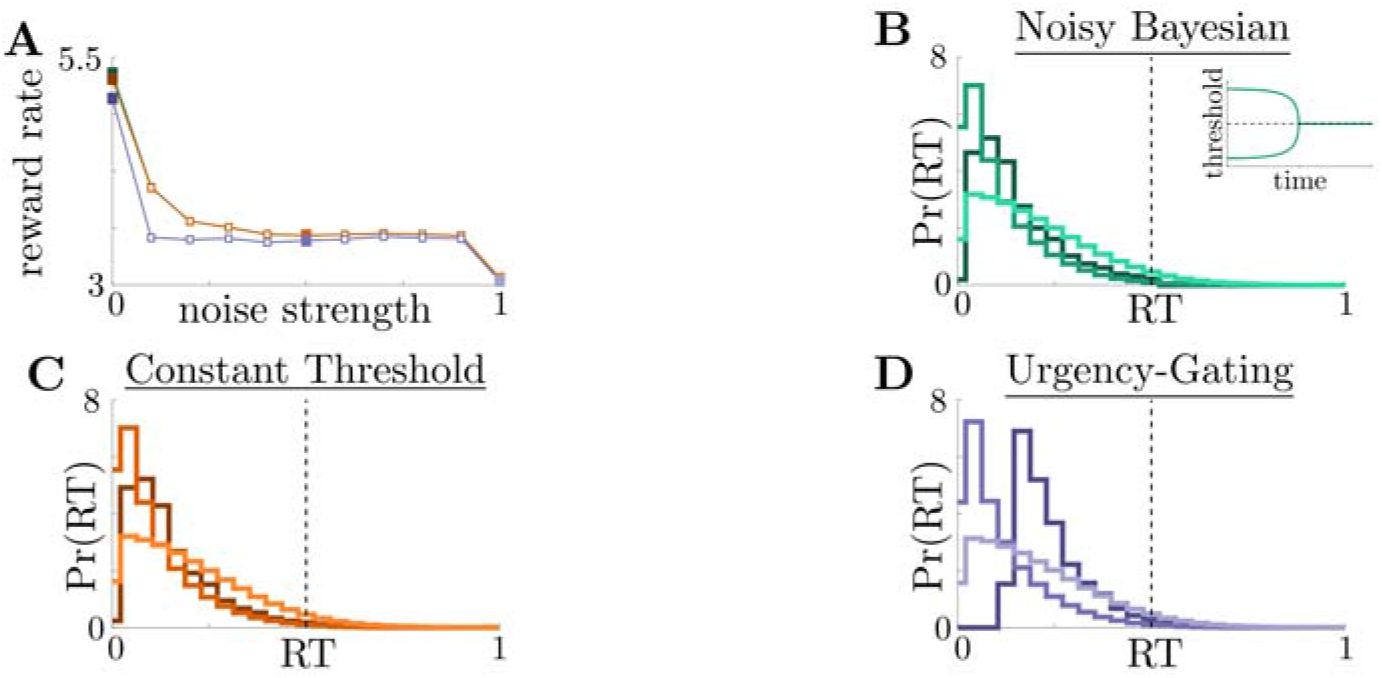
All models perform similarly in high-to-low reward switches. **A**: Reward rates of all models for low-to-high reward changes (dashed line in Figure 4**A**) at different noise strengths. Filled markers correspond to no noise, moderate noise, and high noise strengths. **B,C,D**: Response distribution for (**B**) NB, with inset showing normative thresholds obtained from dynamic programming; (**C**) Const; and (**D**) UGM model in a high-to-low reward environment; several noise strengths, corresponding to filled markers in **A**, are superimposed, with lighter distributions denoting higher noise.

**Figure 4-Figure supplement 3.**
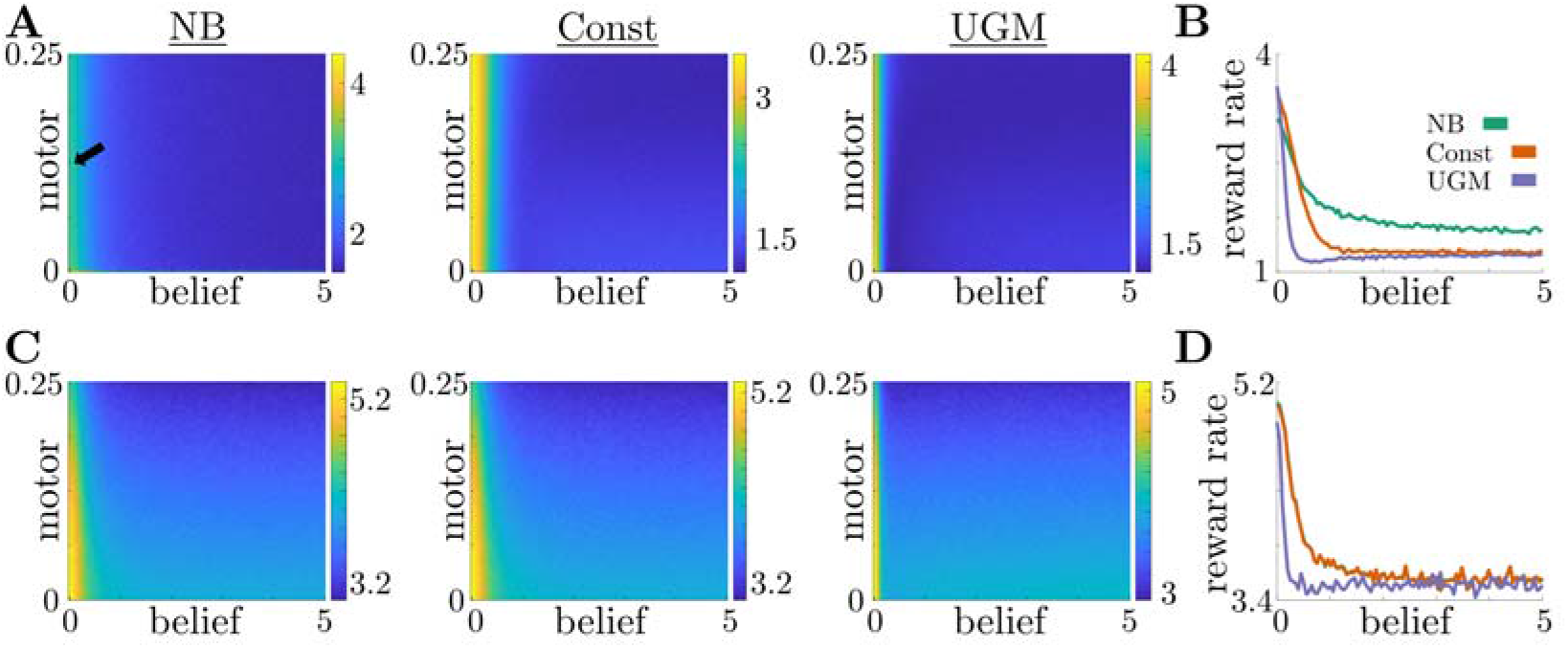
Effects of belief and motor noise on model performance. **A**: Reward rates of NB model (left), Const model (center), and UGM (right) for different values of belief and motor noise strengths in a low-to-high reward switch. Increasing belief noise strength causes performance to decrease substantially, while increasing motor noise strength has little effect on performance. To better visualize performance decreases, we take a slice through the performance surface ata fixed motor noise strength (arrow label in far left panel). **B**: Reward rates of each model for different values of belief noise strength and motor noise strength fixed at 0.125 (arrow label in **A**). **C,D**: Same as **A,B**, but for a high-to-low reward switch.

**Figure 5-Figure supplement 1.**
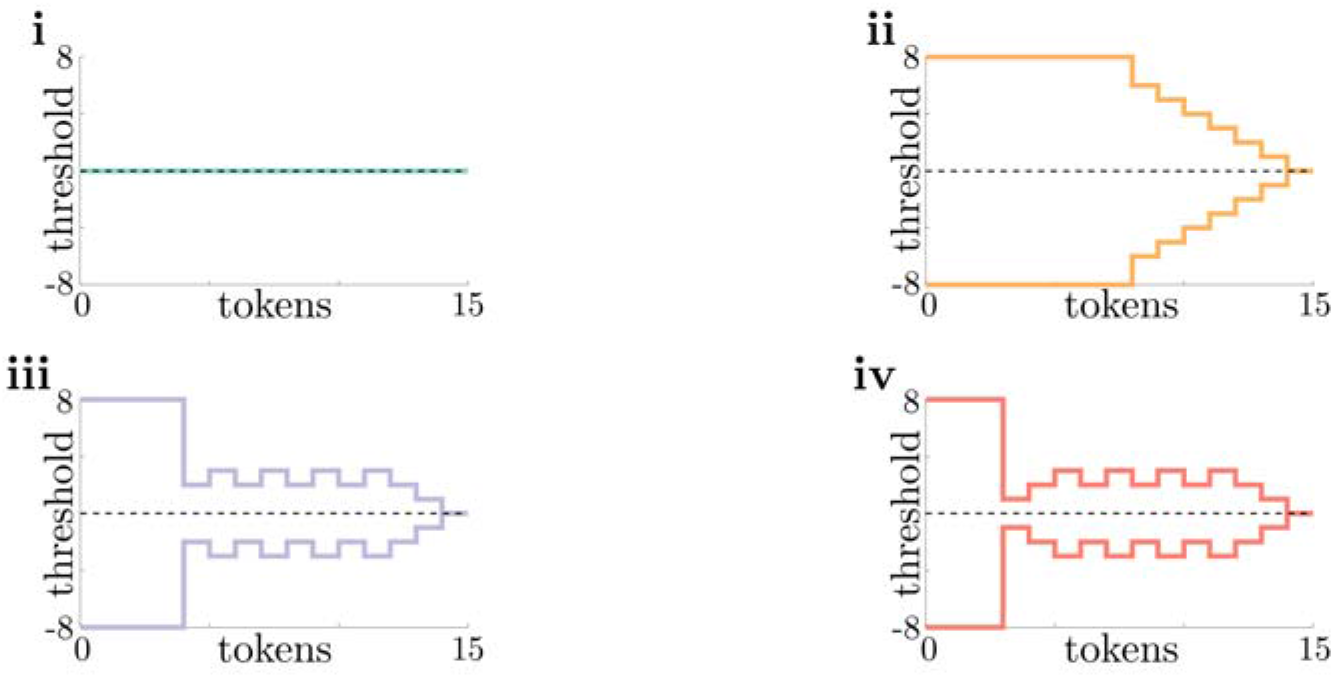
Normative thresholds for tokens task plotted in token lead space. **i-iv**: Same as Figure 5**i-iv**, but plotted in “token lead” space instead of LLR space. Here, thresholds are measured as the number of tokens the top target must be ahead of the bottom target to commit to a decision. In this space, non-monotonicity of thresholds in **iii** and **iv** is more apparent.

**Figure 6-Figure supplement 1.**
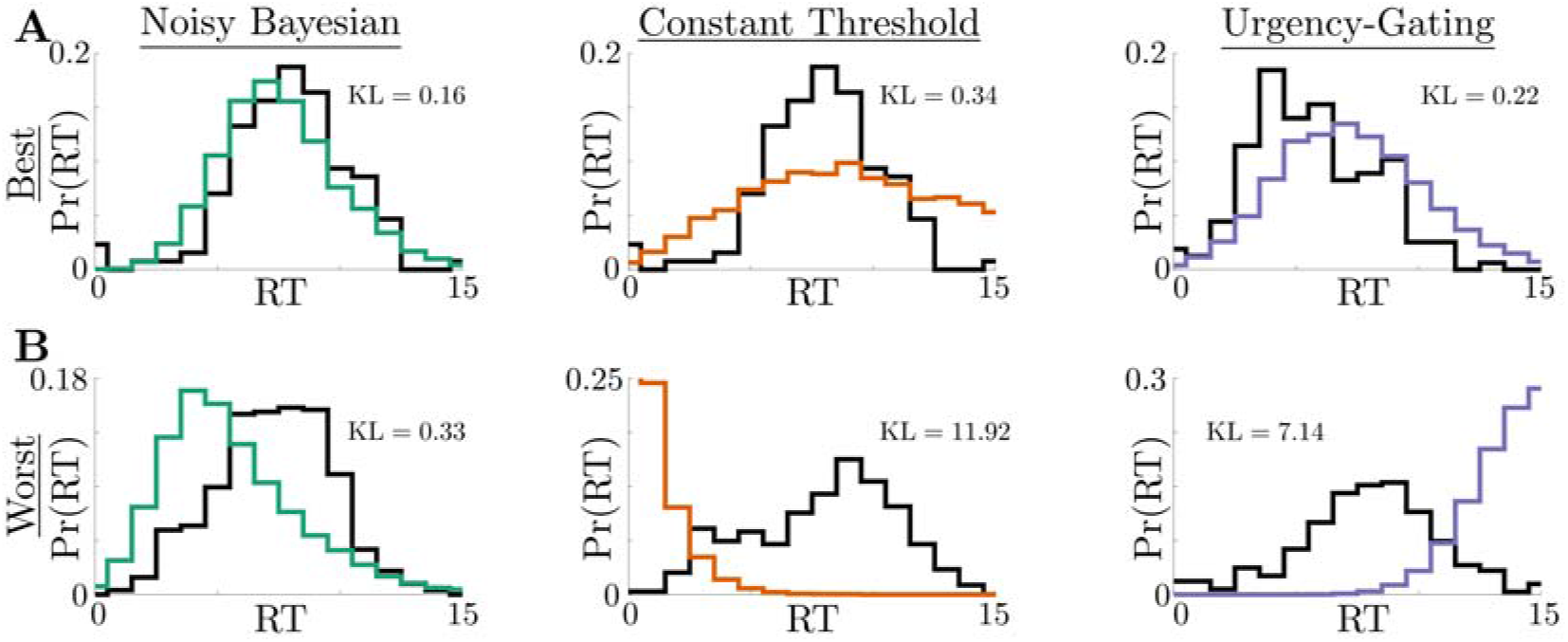
Best- and worst-quality fits for each model. **A**: Best fits, mea-sured using AICc, for the NB model (left), Const model (center), and UGM (right). Black trace shows subject data, and colored trace shows the maximum-likelihood model fit. Each plot shows the Kullback-Leibeler (KL) divergence between the subject data and the fitted response distribution (see Methods and Materials for details). **B**: Same as **A**, but for the worst fits for each model.

## Notes

**Competing interests**: JIG: Senior editor, eLife. The other authors declare that no competing interests exist.

**Funding**: This work was funded by CRCNS/NIH R01-MH-115557. NWB and ZPK were also supported by R01-EB029847-01 and NSF-DMS-1853630. KJ was also supported by NSF DBI-1707400.

https://github.com/nwbarendregt/AdaptNormThresh

### Competing Interest Statement

Joshua Gold: Senior editor, eLife.
The other authors declare that no competing interests exist.

## References

Ashwood ZC, Roy NA, Stone IR, Urai AE, Churchland AK, Pouget A, Pillow JW. Mice alternate between discrete strategies during perceptual decision-making. Nature Neuroscience. 2022; p. 1–12.

Balci F, Simen P, Niyogi R, Saxe A, Hughes JA, Holmes P, Cohen JD. Acquisition of decision making criteria: reward rate ultimately beats accuracy. Attention, Perception, & Psychophysics. 2011; 73(2):640–657.

Barendregt NW, Josić K, Kilpatrick ZP. Analyzing dynamic decision-making models using Chapman-Kolmogorov equations. Journal of computational neuroscience. 2019; 47(2-3):205–222.

Bellman R. Dynamic Programming. Princeton University Press; 1957.

Berger T. Rate-distortion theory. Wiley Encyclopedia of Telecommunications. 2003;.

Bertsekas D. Dynamic programming and optimal control: Volume I, vol. 1. Athena scientific; 2012.

Boehm U, van Maanen L, Evans NJ, Brown SD, Wagenmakers EJ. A theoretical analysis of the reward rate optimality of collapsing decision criteria. Attention, Perception, & Psychophysics. 2020; 82(3):1520–1534.

Bogacz R, Brown E, Moehlis J, Holmes P, Cohen JD. The physics of optimal decision making: a formal analysis of models of performance in two-alternative forced-choice tasks. Psychological review. 2006; 113(4):700.

Bogacz R, Wagenmakers EJ, Forstmann BU, Nieuwenhuis S. The neural basis of the speed-accuracy tradeoff. Trends in neurosciences. 2010; 33(1):10–16.

Brunham K, Anderson D. Model selection and multimodel inference: A practical information-theoretic approach. New York Inc: Springer. 2002;.

Carland MA, Thura D, Cisek P. The urgency-gating model can explain the effects of early evidence. Psychonomic bulletin & review. 2015; 22(6):1830–1838.

Cavanaugh JE. Unifying the derivations for the Akaike and corrected Akaike information criteria. Statistics & Probability Letters. 1997; 33(2):201–208.

Chittka L, Skorupski P, Raine NE. Speed-accuracy tradeoffs in animal decision making. Trends in ecology & evolution. 2009; 24(7):400–407.

Cisek P, Puskas GA, El-Murr S. Decisions in changing conditions: the urgency-gating model. Journal of Neuroscience. 2009; 29(37):11560–11571.

Combes S, Rundle D, Iwasaki J, Crall JD. Linking biomechanics and ecology through predator-prey interactions: flight performance of dragonflies and their prey. Journal of Experimental Biology. 2012; 215(6):903–913.

Crapse TB, Lau H, Basso MA. A role for the superior colliculus in decision criteria. Neuron. 2018; 97(1):181–194.

Drugowitsch J, Notes on Normative Solutions to the Speed-Accuracy Trade-Off in Preceptual Decision-Making; 2015.

Drugowitsch J, Moreno-Bote R, Churchland AK, Shadlen MN, Pouget A. The cost of accumulating evidence in perceptual decision making. Journal of Neuroscience. 2012; 32(11):3612–3628.

Drugowitsch J, Moreno-Bote R, Pouget A. Optimal decision-making with time-varying evidence reliability. In: Advances in neural information processing systems; 2014. p. 748–756.

Drugowitsch J, Moreno-Bote R, Pouget A. Relation between belief and performance in perceptual decision making. PloS one. 2014; 9(5):e96511.

Einfalt LM, Grace EJ, Wahl DH. Effects of simulated light intensity, habitat complexity and forage type on predator-prey interactions in walleye S ander vitreus. Ecology of Freshwater Fish. 2012; 21(4):560–569.

Eissa TL, Gold JI, Josić K, Kilpatrick ZP. Suboptimal human inference inverts the bias-variance trade-off for decisions with asymmetric evidence. bioRxiv. 2021; p. 2020–12.

Evans NJ, Trueblood JS, Holmes WR. A parameter recovery assessment of time-variant models of decision-making. Behavior Research Methods. 2019; p. 1–14.

Faisal AA, Selen LP, Wolpert DM. Noise in the nervous system. Nature reviews neuroscience. 2008; 9(4):292–303.

Glaze CM, Kable JW, Gold JI. Normative evidence accumulation in unpredictable environments. Elife. 2015; 4:e08825.

Glickman M, Moran R, Usher M. Evidence integration and decision confidence are modulated by stimulus consistency. Nature Human Behaviour. 2022; p. 1–12.

Gold JI, Shadlen MN. Banburismus and the brain: decoding the relationship between sensory stimuli, decisions, and reward. Neuron. 2002; 36(2):299–308.

Gold JI, Shadlen MN. The neural basis of decision making. Annu Rev Neurosci. 2007; 30:535–574.

Hanks TD, Kopec CD, Brunton BW, Duan CA, Erlich JC, Brody CD. Distinct relationships of parietal and prefrontal cortices to evidence accumulation. Nature. 2015; 520(7546):220–223.

Jun EJ, Bautista AR, Nunez MD, Allen DC, Tak JH, Alvarez E, Basso MA. Causal role for the primate superior colliculus in the computation of evidence for perceptual decisions. Nature neuroscience. 2021; 24(8):1121–1131.

Kilpatrick ZP, Holmes WR, Eissa TL, Josić K. Optimal models of decision-making in dynamic environments. Current opinion in neurobiology. 2019; 58:54–60.

Louie K, Glimcher PW, Webb R. Adaptive neural coding: from biological to behavioral decision-making. Current opinion in behavioral sciences. 2015; 5:91–99.

Ma WJ, Jazayeri M. Neural coding of uncertainty and probability. Annual review of neuroscience. 2014; 37:205–220.

Mahadevan S. Average reward reinforcement learning: Foundations, algorithms, and empirical results. Machine learning. 1996; 22(1-3):159–195.

Malhotra G, Leslie DS, Ludwig CJ, Bogacz R. Overcoming indecision by changing the decision boundary. Journal of Experimental Psychology: General. 2017; 146(6):776.

Malhotra G, Leslie DS, Ludwig CJ, Bogacz R. Time-varying decision boundaries: insights from optimality analysis. Psychonomic bulletin & review. 2018; p. 1–26.

Palestro JJ, Weichart E, Sederberg PB, Turner BM. Some task demands induce collapsing bounds: Evidence from a behavioral analysis. Psychonomic bulletin & review. 2018; 25(4):1225–1248.

Radillo AE, Veliz-Cuba A, Josić K, Kilpatrick ZP. Performance of normative and approximate evidence accumulation on the dynamic clicks task. Neurons, Behavior, Data analysis, and Theory. 2019; p. 10226.

Ratcliff R. A theory of memory retrieval. Psychological review. 1978; 85(2):59.

Shinn M, Lam NH, Murray JD. A flexible framework for simulating and fitting generalized drift-diffusion models. ELife. 2020; 9:e56938.

Simen P, Contreras D, Buck C, Hu P, Holmes P, Cohen JD. Reward rate optimization in two-alternative decision making: empirical tests of theoretical predictions. Journal of Experimental Psychology: Human Perception and Performance. 2009; 35(6):1865.

Sutton RS, Barto AG, et al. Introduction to reinforcement learning, vol. 135. MIT press Cambridge; 1998.

Tajima S, Drugowitsch J, Patel N, Pouget A. Optimal policy for multi-alternative decisions. Nature neuroscience. 2019; 22(9):1503–1511.

Tajima S, Drugowitsch J, Pouget A. Optimal policy for value-based decision-making. Nature communications. 2016; 7:12400.

Thura D, Beauregard-Racine J, Fradet CW, Cisek P. Decision making by urgency gating: theory and experimental support. Journal of neurophysiology. 2012; 108(11):2912–2930.

Thura D, Cisek P. Modulation of premotor and primary motor cortical activity during volitional adjustments of speed-accuracy trade-offs. Journal of Neuroscience. 2016; 36(3):938–956.

Thura D, Cisek P. The basal ganglia do not select reach targets but control the urgency of commitment. Neuron. 2017; 95(5):1160–1170.

Thura D, Cisek P. Microstimulation of dorsal premotor and primary motor cortex delays the volitional commitment to an action choice. Journal of neurophysiology. 2020; 123(3):927–935.

Thura D, Cos I, Trung J, Cisek P. Context-dependent urgency influences speed-accuracy trade-offs in decision-making and movement execution. Journal of Neuroscience. 2014; 34(49):16442–16454.

Trueblood JS, Heathcote A, Evans NJ, Holmes WR. Urgency, leakage, and the relative nature of information processing in decision-making. Psychological Review. 2021; 128(1):160.

Veliz-Cuba A, Kilpatrick ZP, Josic K. Stochastic models of evidence accumulation in changing environments. SIAM Review. 2016; 58(2):264–289.

Wald A. Sequential tests of statistical hypotheses. The annals of mathematical statistics. 1945; 16(2):117–186.

